# Plant neighborhood shapes diversity and reduces interspecific variation of the phyllosphere microbiome

**DOI:** 10.1101/2021.09.27.462052

**Authors:** Kyle M. Meyer, Robert Porch, Isabella E. Muscettola, Ana Luisa S. Vasconcelos, Julia K. Sherman, C. Jessica E. Metcalf, Steven E. Lindow, Britt Koskella

## Abstract

Microbial communities associated with plant leaf surfaces (i.e. the phyllosphere) are increasingly recognized for their role in plant health. While accumulating evidence suggests a role for host filtering of its microbiota, far less is known about how community composition is shaped by dispersal, including from neighboring plants. We experimentally manipulated the local plant neighborhood within which tomato, pepper, or bean plants were grown in a three-month field trial. Focal plants were grown in the presence of con- or hetero-specific neighbors (or no neighbors) in a fully factorial combination. At 30-day intervals, focal plants were harvested and replaced with a new age- and species-matched cohort while allowing neighborhood plants to continue growing. 16S community profiling revealed that the strength of host filtering effects (i.e. interspecific differences in composition) decreased over time. In contrast, the strength of neighborhood effects increased over time, suggesting dispersal from neighboring plants becomes more important as neighboring plant biomass increases. We next implemented a cross-inoculation study in the greenhouse using inoculum generated from the field plants to directly test host filtering of microbiomes while controlling for directionality and source of dispersal. This experiment further demonstrated that focal host species, the host from which the microbiome came, and in one case the donor hosts’ neighbors, contribute to variation in phyllosphere bacterial composition. Overall, our results suggest that local dispersal is a key factor in phyllosphere assembly, and that demographic factors such as nearby neighbor identity and biomass or age are important determinants of phyllosphere microbiome diversity.

## Introduction

Plant leaf surfaces, commonly termed the phyllosphere, harbor a wide diversity of microorganisms ^1^. These endophytic and epiphytic communities can influence plant health and fitness through a variety of means, including protection against pathogens ^2,3^, plant growth promotion ^4^, primary productivity enhancement ^5^, protection against abiotic conditions including frost ^6^, and fixation of atmospheric nitrogen (N) ^7^. Plant hosts can exert some control over the abundance and composition of their microbiome members by virtue of the differing chemical and physical features of resources provided on their surfaces ^8^, but also through immune activity, molecular signaling, and barrier formation ^9–15^. This filtering effect can give rise to predictable differences in microbiome composition among hosts ^10,16–18^, a phenomenon referred to as species identity (or genotype) effects. Evidence for such effects comes from phylogenetic clustering of associated microbial taxa ^19,20^, deviation from null or neutral expectations ^21,22^, changes consequent to host genetic manipulation ^10^, or compositional differences explained by species or genotype as a factor ^16–18,20,23^. While species identity effects suggest the importance of host control over microbiota, such effects are often weak or variable when tested in broader environmental or ecological contexts ^24,25^. This raises the question of whether and how host effects can be swamped by environmental factors in shaping the microbiome.

The neighboring plant community constitutes a major component of a plant’s environmental and ecological context. Neighborhood effects, also known as associational effects, have been extensively studied for pathogen and herbivore transmission ^26–28^, revealing patterns of transmission that relate to the nearest conspecific neighbor (i.e. conspecific negative density dependence) ^29–32^ as well as species frequency-dependent patterns of host fitness ^33–35^. Much less effort has focused on the role of neighborhood effects for non-pathogenic plant-associated microorganisms ^36^. Given the prominent role of aerial transmission in shaping phyllosphere microbial communities ^23,37–39^, both neighbor identity and proximity are likely to be important factors shaping epiphytic microbial communities. Moreover, it has been shown both theoretically ^40^ and empirically ^41^ that in the presence of high dispersal rates, community members can persist even in the face of strong selection against them (e.g. as a result of plant filtering effects), a phenomenon termed mass effects. As such, differences in microbiota composition that arise between species could be diminished when inter-host dispersal is high. Indeed this has been shown in zebrafish, where differences in bacterial community composition among host variants were dramatically reduced when inter-host dispersal was allowed ^42^.

Recent observational research in tree communities has revealed detectable neighborhood effects on epiphytic communities ^38^, but many open questions remain. For instance, it is unclear whether neighborhood effects are general and causative, a crucial gap in knowledge if such effects are to be incorporated into agricultural practice. It is also unclear what role neighbor or focal plant identity and age/biomass play in microbiome assembly. Lastly, although host filtering and microbial dispersal are intimately intertwined, the relative impacts of each in shaping microbiome differentiation among species has not been described. We address these knowledge gaps using field- and greenhouse-based experiments involving tomato, pepper, and bean host plants. By manipulating both the focal plant species identity and the local neighborhood composition in the field, we were able to directly test the relative importance of plant host filtering versus local dispersal sources in shaping microbiome composition on leaves. We then performed a controlled cross-inoculation study in the greenhouse to directly examine the effects of host filtering and inoculum source on microbiome assembly of the three plant species involved. We hypothesized that: 1) neighborhood effects would increase as neighboring plants increase in age and biomass; 2) that neighborhood effects would depend on both neighbor and focal plant identity due to host filtering effects; 3) that host species effects would be diminished in the presence of neighboring plants; and 4) that experimental transplantation of microbiomes across hosts would result in compositional change as a result of host filtering, but that there would remain a detectable signal of past host.

## Methods

### Experimental Design: Neighborhood study

To test for the relative influence of host species and neighborhood effects on foliar microbial communities, we implemented a fully factorial, randomized block design at the Oxford Tract, a research farm near the University of California, Berkeley. The study included three plant species: tomato (Solanum lycopersicum var. Moneymaker), pepper (Capsicum anuum var. Early Cal Wonder), and bean (Phaseolus vulgaris var. Bush Blue Lake 274). Plant neighborhoods were established in which a single 5 week old tomato or pepper plant, or a cluster of 6, 2-week old beans, was planted as the focal individual in the middle of a circle of eight neighborhood plants, each planted 0.61 m from the focal plant, with fully reciprocal combinations of focal and neighborhood plants (Fig. 1A) in a randomized complete block design with 6 replicates. Neighborhood plots were established in 3 zones (2 blocks per zone) spaced 1.22m apart separated by at least 0.91m from any other plants at the experimental site to minimize edge effects. Each neighborhood plot was 1.22 x 1.22 m, and separated by 1.22 m from adjacent plots. No neighbor control plots, in which focal plants had no neighbors encircling them, were included for each species. Thus, the experiment contained 9 neighborhood comparisons and 3 no neighbor comparisons. Two weeks prior to planting, soil was tilled for weed management and drip lines were installed underneath plastic sheeting to provide irrigation. The plastic sheeting prevented the growth of weeds and minimized dispersal of soil onto plants. Plants were planted through small holes made through the plastic sheeting. Individual tomato and pepper plants were propagated in a greenhouse for 5 weeks, to a height of about 20 cm before transplantation into the field. Beans, 6 seeds per pot, were grown for 20 days to a height of 20 cm before transplantation. All greenhouse plants were watered using drip irrigation to minimize the wetting of the leaves, and thus the development of large epiphytic bacterial community sizes. All plants, including focal individuals and neighbors, were transplanted in the field June 1, 2019.

**Fig. 1:**
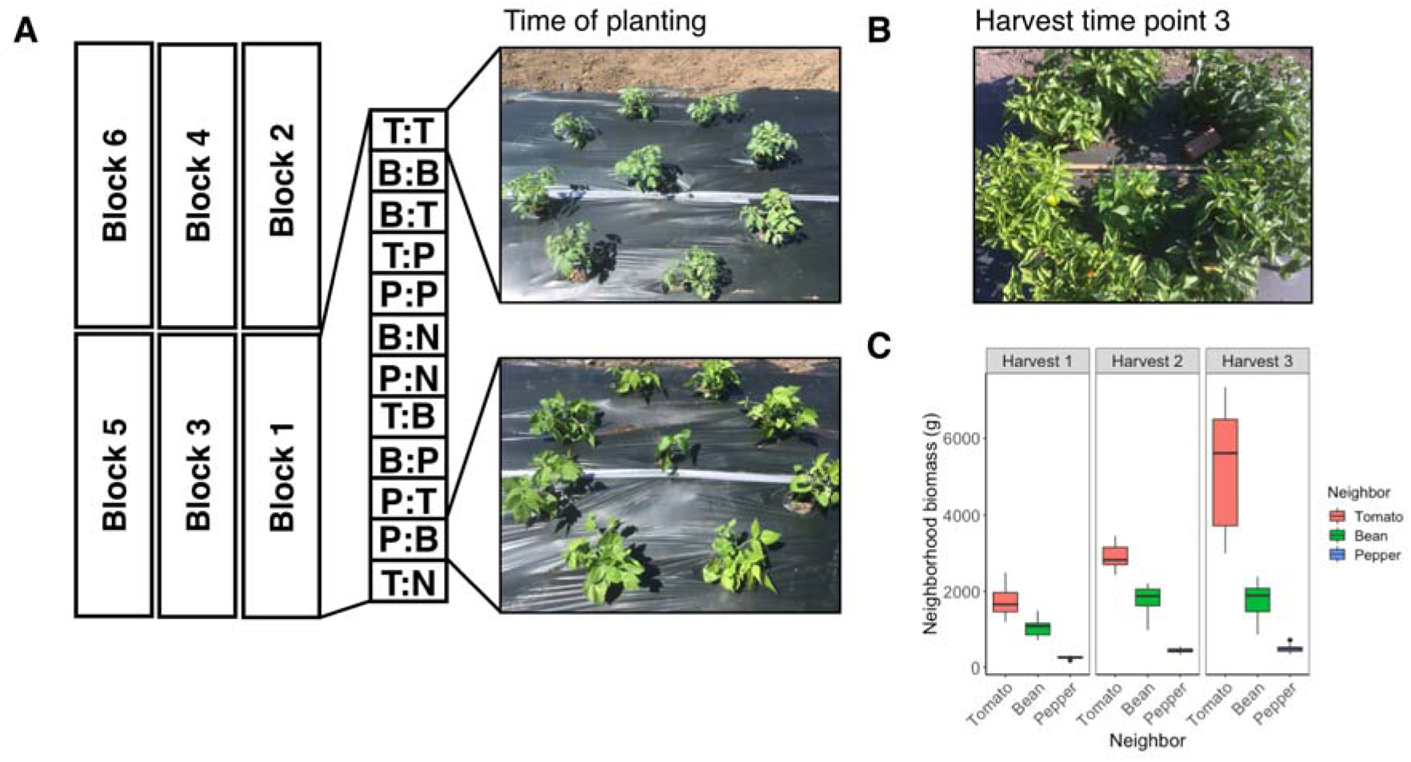
Experimental design of the field trial. A) Experimental neighborhoods were constructed by planting a focal plant in the center of a ring of neighbor plants, with fully factorial combinations of focal and neighbor plant species. Each block contained 12 comparisons, with focal and neighbor plant abbreviated (T=tomato, P=pepper, B=bean, and N=no neighbor), respectively. Focal plants were harvested and replaced each month, while the neighborhoods were left to continue growing. Shown are a tomato focal plant with tomato neighbors (top) and a pepper focal plant with bean neighbors (bottom) at the time of initial planting. B) A pepper neighborhood surrounding a bean focal plant at the time of harvest number 3. C) The biomass of neighborhood plants (g) increased to varying degrees with time for each plant species by harvests 1, 2, and 3.

Focal plants were harvested and replaced at 30-day intervals (3 times of establishment) while the neighborhoods were retained and continued to grow throughout the study (Fig. 1B, C). Focal plants were all the same age upon planting as the original cohorts to allow for direct comparisons across cohorts. Thus, bacterial community composition of the focal plants was assessed on 3 separate occasions for each of 72 focal plants in the neighborhoods (totaling 216 focal samples). Only one focal plant (pepper with no neighbors) died prior to sampling. Additionally, at each round of planting, bacterial community composition was assessed on 5 plants of each species at the time of transplantation to identify taxa that had established on plants in the greenhouse.

### Neighborhood Plant Attribute Measurements

Several attributes of neighborhood plants were measured before each monthly harvest of focal plants in order to determine how neighbors might impact phyllosphere communities of the focal plant. These attributes included: average neighbor height, distance of the focal plants to the nearest neighbor, the number of neighbors touching the focal plant (if any), the total number of flowers on the neighborhood plants, and whether the neighborhood plants had signs of herbivory, infestation, or disease (yes or no). Further, the biomass of the neighborhood was estimated without harvesting the plants by fitting a linear model relating the height and weight of the focal plants, and extrapolating to that of the neighbor plant weights based on their height. Separate linear models were fit for each plant species.

### Sample processing

Immediately before harvesting the focal plants, their height was measured. Focal plants were then excised at their base using ethanol-sterilized scissors, transferred to 1-gallon (3.79 L) sterile plastic bags, and transported in a chilled cooler to the laboratory. Plant weight was recorded and then plants were subsampled, collected, and re-weighed. Subsampling was necessary to reduce biomass differences among samples and to enable more efficient collection of epiphytic bacteria by sonication. Foliar bacteria were collected from plant subsamples (range 3.84 – 621.14 g, median 40.84 g) by adding 180 ml of sterile 10mM MgCl_2_ to the sample bags and sonicating for 10 minutes in a sonicating water bath (Branson model 5800). Leaf wash was then filtered through an autoclaved coffee filter and distributed to four 50 ml conical tubes, which were then centrifuged at 4000 rcf at 10° C for 10 minutes to pellet microbial cells. The supernatant was then decanted from each tube and the pellets resuspended in 1.8 ml King’s broth (KB). 600 μl of the resuspended pellet was frozen at −80° C for subsequent DNA extraction, while the remaining two 600 μl aliquots were each mixed with 400 ul 1:1 KB:Glycerol and frozen at −80° C for subsequent experimentation.

### Experimental Design – Follow-up transplant study

To further test the importance of host filtering, inoculum source, and dispersal history, we conducted a follow-up greenhouse study in which bacterial communities recovered from field plants at harvest time point 2 were reciprocally inoculated onto these same species under controlled conditions. Cryopreserved phyllosphere communities from a single focal field plant were transferred to either the same plant species from which they were isolated, or onto the plant species that previously had surrounded that focal plant when it was in the field. For instance, the microbiome from a tomato that was surrounded by beans was applied equally to a tomato and a bean plant. This was done for all combinations. Experimental blocks from the field trial were treated as experimental blocks in the greenhouse trial, using blocks 2-6 from the field (5 replicates per treatment). We deliberately did not equalize inoculum density, as we anticipated that bacterial abundances would vary according to plant species and thereby constitute an important component of species identity effects. The biomass of every donor plant, however, was recorded for downstream statistical analysis. Additionally, for each plant species we included 5 replicate blank inoculum controls in which the same volume of sterile 10 mM MgCl_2_ that was used to resuspend inoculum was sprayed onto plants. We further included replicate heat-killed controls, in which field-derived leaf wash was autoclaved for 40 minutes before being applied to each of three plant hosts, in the same manner as the experimental inocula.

Inocula were prepared by thawing the freezer stock, centrifuging at 4000 rcf and 10° C for 10 minutes to pellet cells, decanting the supernatant, and re-suspending cells in 7 ml 10 mM MgCl_2_ then splitting in half to make two 3.5 mL inocula. Twenty-two samples were inoculated each day (block) such that each block contained every comparison, and this was repeated for 5 days. Inocula were sprayed onto the adaxial (top) and abaxial (bottom) sides of leaves using ethanol- and UV-sterilized misting caps. After inoculation, the moist, sprayed plants were placed in a chamber maintaining ca. 100% relative humidity for 20 hours in order to maintain leaf moistness, thus encouraging microbial growth, before being transferred to a greenhouse. After 7 days, the plants were returned to the humid chamber for 20 hours immediately before harvest in order to facilitate further microbial multiplication on leaves and thus allow for maximal host filtering. Plants were then harvested and processed as in the field study.

### DNA extraction, PCR, Library Preparation, and Sequencing

One sixth of the total leaf surface microbial extraction per plant was used for DNA extraction with DNeasy Powersoil Kits (Qiagen). Sample order was randomized to avoid batch effects, and a blank (no sample) control was included in every round of DNA extraction. DNA concentration of each sample was quantified using the Qubit dsDNA HS Assay Kit. 10 μl of sample DNA was used as template and PCR amplified for 35 cycles at the University of California - Davis Host Microbe Systems Biology Core using the 799F (5’ – AACMGGATTAGATACCCKG – 3’) - 1193R (5’ – ACGTCATCCCCACCTTCC – 3’) primer combination, which targets the V5-V7 region of the 16S rRNA gene, and was designed to minimize chloroplast amplification ^43,44^. To further minimize host mitochondrial and chloroplast amplification, peptide nucleic acid (PNA) clamps were added to each reaction ^45^. Resulting amplicons were diluted 8:1 and were further amplified for 9 cycles to add sample-specific barcodes, then quantified using Qubit, pooled in equal amounts, cleaned with magnetic beads and size selected via electrophoresis on a Pippin Prep gel (Sage Science, USA). The resultant library was then sequenced on the Ilumina MiSeq (paired-end 300) platform.

### Sequence Processing

Amplicon sequences were processed using the DADA2 pipeline ^46^ implemented in the R statistical environment ^47^, including the packages ShortRead ^48^, Biostrings ^49^, and Phyloseq ^50^. Forward and reverse reads were truncated at 260 and 160 bp, respectively, and quality filtered using the function ‘filterAndTrim’ with default settings (i.e. maxN=0, maxEE=c(2,2), and truncQ=2). Error rates for forward and reverse reads were determined using the ‘learnErrors’ function, and then applied to remove sequencing errors from reads and assign them to amplicon sequence variants (ASVs) using the ‘dada’ function. Filtered paired reads were merged using the function ‘mergePairs’ and then converted into a sequence table using the ‘makeSequenceTable’ function. Chimeric sequences were removed from the sequence table using the function ‘removeBimeraDeNovo’ (method = consensus). Taxonomy was assigned to the remaining sequences using the ‘assignTaxonomy’ function, which implements the RDP Naïve Bayesian Classifier algorithm with kmer size 8 and 100 bootstrap replicates ^51^. We used the Silva SSU taxonomic training dataset (version 138) formatted for DADA2 ^52^. Chloroplast and mitochondrial sequences were filtered from the ASV table by removing any ASVs with a taxonomic assignment of ‘Chloroplast’ at the Order level or ‘Mitochondria’ at the Family level, respectively. Lastly, we applied the ‘isContaminant’ function (method = prevalence) from the package ‘decontam’ ^53^ to our samples using our blank (no sample) DNA extractions to identify and remove putative contaminants introduced during DNA extraction.

### Bacterial Quantification using Droplet Digital PCR (ddPCR)

In order to estimate foliar bacterial abundances of each plant sample, droplet digital PCR (ddPCR) using the Bio-Rad QX200 system on bacterial DNA extracted from leaf wash. Comprehensive ddPCR methods are described elsewhere ^54^, but briefly, we targeted the V5-V7 region of the 16S rRNA gene in sample DNA using the chloroplast-excluding 799F (5’ – AACMGGATTAGATACCCKG – 3’) – 1389R (5’ – ACGGGCGGTGTGTRC – 3’) primer combination. 5 μl of 1:10 diluted DNA template were combined with 11 μl of 2X EvaGreen Supermix (Bio-Rad, USA) and 0.22 μl of each primer, and 5.56 μl of molecular grade water to a total volume of 20 μl. Reaction mixes were then loaded into the QX200 droplet generator with 70 ul of droplet generation oil, then transferred to a PCR plate. 39 cycles of PCR were performed under the following conditions: 95°C for 10 minutes, 95°C for 30 seconds, 55°C for 30 seconds, 72°C for 2 minutes, with steps 2-4 repeated 39 times, 4°C for 5 minutes, and 90°C for 5 minutes. EvaGreen signal was measured on the QX200 droplet reader, cutoff thresholds were set for each column based on background fluorescence in no template controls, and concentrations were determined using the associated QuantaSoft software. Abundances are reported as 16S rRNA copies per g plant material as well as estimates of 16S copies per individual plant by taking into account the proportion of the total plant sample that was used for sample processing.

### Statistical Analysis

All statistical analyses were performed using R version 4.0.3 ^47^. Community matrices were rarefied to 6400 counts per sample ten times and averaged in order to account for differences in sampling extent across samples. Bray Curtis bacterial community dissimilarities were calculated between samples using the ‘vegdist’ function in the vegan package in R ^55^. Community structure differences among host species identity, neighbor species identity, and experimental block were assessed using a PERMANOVA on Bray-Curtis distances using the ‘adonis’ function (also in the vegan package), which performs a sequential test of terms and uses the algorithm presented in ^56^. To assess the change in the relative strength of these factors through time, the PERMANOVA was performed for each of the three harvesting time points separately. Since not all samples successfully sequenced, generating slight differences in sample numbers among harvests, we adjusted R^2^ values to take into account sample numbers and degrees of freedom using the ‘RsquareAdj’ function in the vegan package. Indicator taxa analysis was performed using the ‘multipatt’ function in the indicspecies package ^57^.

In order to assess the unique contribution of plant species identity for each neighborhood type, we used variation partitioning on Hellinger-transformed community matrices to partition out the effects of space. A geographic distance matrix was calculated for all experimental plots based on plot GPS coordinates using the program Geographic Distance Matrix Generator ^58^, and then pairwise distances were decomposed into principal coordinates using the ‘pcnm’ function in the vegan package. For each combination of plants, significant principal coordinates were selected using forward and backward (direction = “both”) model selection with the ‘ordistep’ function in the vegan package. The unique contribution of host species identity was then calculated after partitioning out the significant spatial PCNMs that were selected using the ‘varpart’ function in vegan. We further assessed the contribution of other host factors including plant height and weight by performing the above-mentioned model selection and variation partitioning. The statistical significance of variation fractions was then tested by performing redundancy analysis ordination (RDA), and declaring the non-focal factors as conditions.

Neutral modeling of phyllosphere communities was performed using the ‘fit_sncm’ function in the package reltools ^59^. This package fits the neutral model from ^60^, as implemented by ^21^. In order to assess phylogenetic patterns in the phyllosphere communities, we constructed a phylogenetic tree of all ASVs with greater than 20 counts in the community matrix, which included 7949 ASVs. Sequences were aligned using the ‘AlignSeqs’ function in the DECIPHER package ^61^ using default settings. Next, pairwise distances between sequences were calculated using the ‘dist.ml’ function in the phangorn package version 2.5.5 ^62^. These distances were then used to construct a neighbor-joining tree using the ‘NJ’ function in phangorn. Lastly, the neighbor-joining tree was used as a starting point to create a generalized time-reversible with gamma rate variation (GTR+G+I) maximum likelihood tree using the ‘pml’, ‘update’, and ‘optim.pml’ functions in the phangorn package. Lastly, we calculated the mean pairwise distance (MPD) of taxa in each sample and compared to the MPD of a null model to calculate the standardized effect size (SES) using the ‘ses.mpd’ function in the picante package ^63^. We used a community randomization null model (null.model = species.pool, iterations=999) whereby a randomized community matrix is constructed by drawing from the total species pool with equal probability. Using such a procedure, a Z-score (the SES of MPD versus the null community) below zero can be interpreted as phylogenetically clustered whereby taxa co-occurring in a sample are more closely related than the same number of taxa drawn at random from the species pool. By contrast, samples above zero are interpreted as phylogenetically overdispersed, i.e. phylogenetic distance among co-occurring taxa is greater than the above-stated null expectation.

For univariate data such as ASV-level richness, MPD SES, and ddPCR-based abundance data, a three-way ANOVA was fit to test for significant effects of host, neighbor, and harvest time point, with interactions therein. The appropriateness of this procedure was verified by checking for a normal distribution of residuals on the model.

## Results

### Experimental manipulation of plant neighborhood in the field

We first compared the phyllosphere microbiome of plants that were surrounded by no neighbors, conspecific (same species) neighbors, or heterospecific (different species) neighbors. Because the field trial was conducted over the course of 3 months, with focal plants being replaced with a plant of the same species but at the original age of planting each month, we were also able to compare neighborhood age/biomass effects on microbiome assembly. After processing, 175 of the 216 focal plant samples from the field yielded high quality sequencing reads (Tomato n = 63, Pepper n = 52, Bean n = 60). Of these 175, 64 were from harvest 1, 58 were from harvest 2, and 53 were from harvest 3. The 41 excluded samples each had less than 24 total reads and thus failed either to amplify or to yield sequences. The greenhouse control plants that were included to assess the bacterial communities established prior to transplantation into the field yielded very few reads (28 of the 45 samples had less than 50 reads), indicating that bacterial colonization prior to transplantation was minimal. Of the 17 greenhouse control samples with detectable sequences, communities were dominated by Enterobacterales, Corynebacterales, Burkholderiales, and Pseudomonadales.

The field trial dataset contained 5,414,393 observations of 19,818 ASVs, 13,455 of which had >10 occurrences, and 2,253 of which had >100 occurrences. Within-host ASV-level richness ranged from 22 to 769 ASVs across all treatments and hosts. Richness levels varied significantly by harvest time (F_2,174_ = 24.21, p < 0.001), declining throughout the season, and varying by host identity (F_2,174_ = 4.96, p < 0.01), with beans harboring a greater richness, especially at the first harvest. Neighborhood did not impact bacterial richness, however the total number of flowers on the neighborhood plant species at the time of focal plant harvest was positively correlated with bacterial richness on the focal plants (R^2^ = 0.044, p = 0.01). Similar qualitative trends were observed for Shannon diversity, except that host plant weight and height were also positively correlated with diversity (R^2^_adj_ = 0.022, p = 0.03 and R^2^_adj_ = 0.044, p < 0.01, respectively).

Bacterial abundance per plant varied significantly by host identity (F_2,174_ = 27.35, p < 0.001), neighborhood (F_3,174_ = 7.96, p < 0.001), and harvest time point (F_2,174_= 3.85, p = 0.051, Fig. 2A), with no significant interactions among variables. Tomato and bean plants tended to harbor higher bacterial abundances than pepper plants (p < 0.001). As expected, abundance per plant was positively correlated with plant weight (R^2^ = 0.187, p < 0.001) and plant height (R^2^ = 0.117, p < 0.001). If we normalize bacterial abundances by the plant material weight used for processing, we see weaker effects of host identity (F_2,174_ = 3.24, p = 0.041) and neighbor (F_3,174_ = 2.17, p = 0.094), but a stronger effect of harvest time point (F_2,174_ = 5.15, p = 0.024). Several neighborhood attributes had interesting associations with bacterial abundance on focal hosts. Specifically, estimated neighborhood biomass (R^2^ = 0.146, p < 0.001), average neighbor height (R^2^ = 0.123, p < 0.001), and total number of flowers (R^2^ = 0.022, p = 0.038) were all negatively correlated with bacterial abundance on focal plants. Lastly, bacterial abundance per focal plant was negatively associated with community richness (R^2^ = 0.016, p = 0.05) and Shannon diversity (R^2^ = 0.03, p < 0.01).

**Fig. 2:**
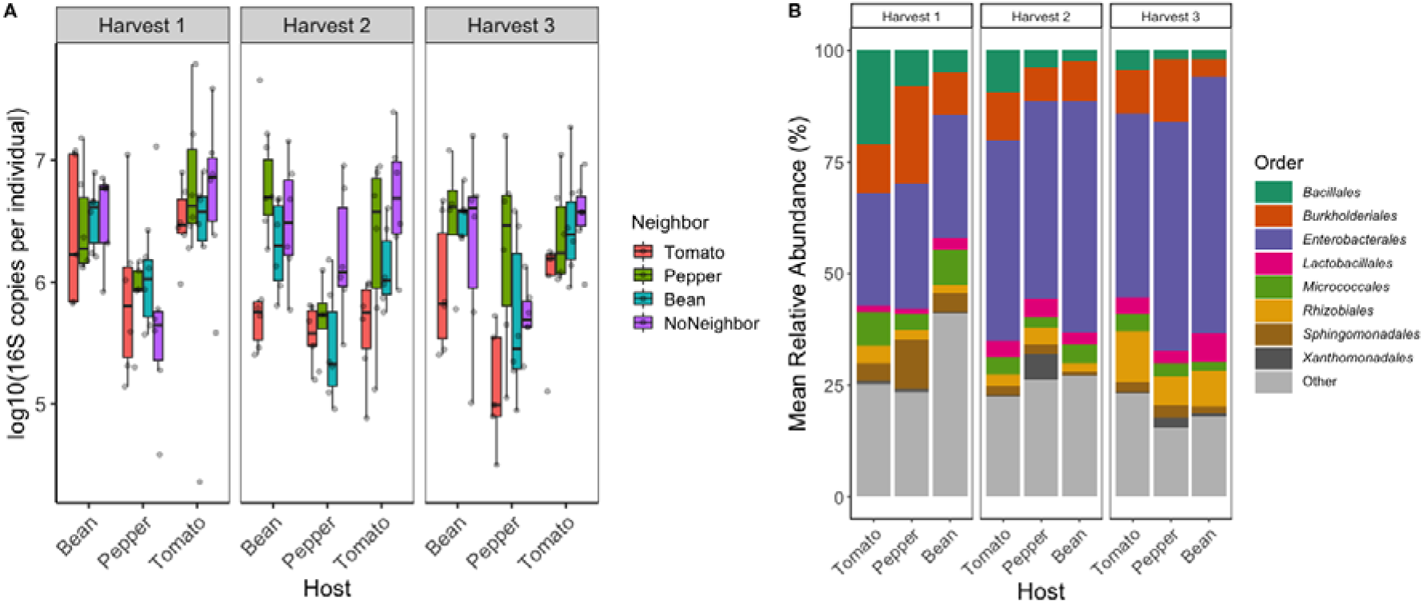
Bacterial abundance and composition vary across host species and harvest time. A) The abundance (log_10_ 16S rRNA gene copies measured using ddPCR of leaf washes, y-axis) for individual focal plant species (x-axis) surrounded by different neighbor plant species (box color) and at different successive harvest times (panels 1, 2, and 3). B) Relative abundance of the 9 most abundant bacterial orders distinguished by host species and harvest time. All other less abundant or ambiguously assigned orders are grouped under ‘Other’.

Overall, phyllosphere communities were dominated by the phyla Proteobacteria, Firmicutes, and Actinobacteriota. The most abundant bacterial orders were the Bacillales, Burkholderiales, Enterobacterales, Lactobacillales, Micrococcales, Rhizobiales, Sphingomonadales, and Xanthomonadales (Fig. 2B). Bray Curtis dissimilarities among samples were driven by harvest time point (R^2^ = 0.063, p = 0.003), host species (R^2^ = 0.055, p = 0.003), neighbor (R^2^ = 0.023, p = 0.048), and block (R^2^ = 0.036, p = 0.06), with a significant interaction between host and harvest (R^2^ = 0.034, p = 0.033), and a trending interaction between neighbor and harvest (R^2^ = 0.041, p = 0.072).

#### The effects of host identity on bacterial community composition decrease through time while neighborhood effects increase through time and vary by host identity

We next examined the relative influence of host species identity and neighborhood on focal plant microbiome structure at each time point during the field experiment by performing a PERMANOVA on Bray Curtis dissimilarities using host species identity (i.e. tomato, pepper, or bean), neighborhood (i.e. tomato, pepper, bean, or no neighbor), and experimental block (1-6) as independent variables. The effect of host identity was significant, but diminished in size over the three time points (Harvest 1: Adj. R^2^ = 0.096, p < 0.001; Harvest 2: Adj. R^2^ = 0.068, p < 0.001; Harvest 3: Adj. R^2^ = 0.027, p < 0.001, Fig. 3A, see Table 1 for pre-adjusted R^2^ values). In contrast, the effect of neighborhood status was initially not statistically significant, but increased in size over the three time points (Harvest 1: Adj. R^2^ = −0.001, p = 0.242; Harvest 2: Adj. R^2^ = 0.017, p < 0.01; Harvest 3: Adj. R^2^ = 0.032, p < 0.001, Table 1, Fig 3A). Block effects tended to decrease throughout the experiment (Harvest 1: Adj. R^2^ = 0.011, p = 0.021, Harvest 2: Adj. R^2^ = 0.009, p = 0.028, Harvest 3: Adj. R^2^ = 0.009, p = 0.07, Fig. 3A). No significant interactions among variables were observed at individual harvests.

**Fig. 3:**
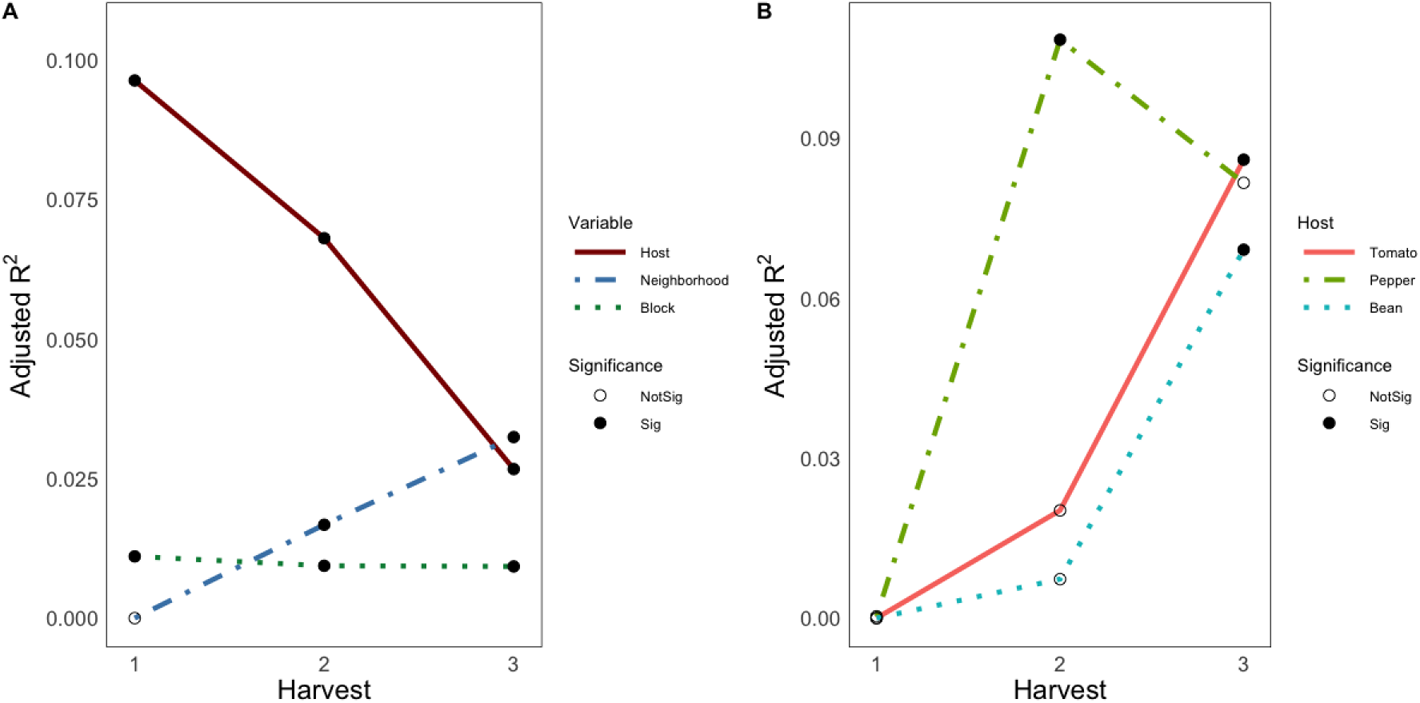
The effects of host identity on bacterial community composition decrease through time while neighborhood effects increase and vary by host plant species. A) Adjusted R^2^ values (y-axis) are the result of PERMANOVA analyses on Bray Curtis dissimilarities for each harvest, accounting for sample number and degrees of freedom from slight differences in sample number. See Table 1 for pre-adjusted R^2^ values. The effect of host identity (solid maroon line), the effect of neighborhood (blue dot-dash line), and the effect of experimental block (green dotted line) are shown. Harvest time point is shown on x-axis. Filled circles indicate statistical significance (p < 0.05), while open circles represent statistically insignificant effects (p > 0.05). B) Host plant species experience neighborhood effects on phyllosphere bacterial communities differently through time. Adjusted R^2^ values (y-axis) and harvest time point (x-axis) are as described for plot A. Tomato hosts (solid red line), pepper hosts (light green dot-dash line), and bean hosts (light blue dotted line) are shown. Filled circles indicate statistical significance (p < 0.05), while open circles represent statistically insignificant effects (p > 0.05).

**Table 1:** Results of a PERMANOVA on phyllosphere bacterial community Bray Curtis dissimilarities for harvest time points 1, 2, and 3 in the field trial. Variables tested include: host species identity (tomato, pepper, or bean), neighborhood (tomato, pepper, bean, no neighbor), and experimental block (1 through 6). R^2^ values represent the fit of the model and adjusted R^2^ values have been adjusted based on sample numbers of degrees of freedom to render values comparable across harvest time points.

| Harvest 1 |  |  |  |  |  |  |
| --- | --- | --- | --- | --- | --- | --- |
|  | *df* | *Pseudo-f* | *R*<sup>2</sup> | *Adj. R*<sup>2</sup> | *P* | *Significance* |
| Host Species | 2 | 4.49 | **0.125** | **0.096** | **0.001** | *** |
| Neighborhood | 3 | 1.11 | 0.046 | -0.001 | 0.242 | NS |
| Block | 5 | 1.28 | 0.089 | **0.011** | **0.021** | * |
| Harvest 2 |  |  |  |  |  |  |
|  | *df* | *Pseudo-f* | *R*<sup>2</sup> | *Adj. R*<sup>2</sup> | *P* | *Significance* |
| Host Species | 2 | 3.22 | **0.1** | **0.068** | **0.001** | *** |
| Neighborhood | 3 | 1.46 | **0.069** | **0.017** | **0.001** | *** |
| Block | 5 | 1.23 | **0.096** | **0.009** | **0.028** | * |
| Harvest 3 |  |  |  |  |  |  |
|  | *df* | *Pseudo-f* | *R*<sup>2</sup> | *Adj. R*<sup>2</sup> | *P* | *Significance* |
| Host Species | 2 | 1.81 | **0.064** | **0.027** | **0.001** | *** |
| Neighborhood | 3 | 1.66 | **0.088** | **0.032** | **0.001** | *** |
| Block | 5 | 1.18 | 0.104 | 0.009 | 0.07 | . |

By excluding the ‘no neighbor’ controls, we then tested for an effect of neighbor identity by treating neighbor type (i.e. tomato, bean, or pepper) as an independent variable. On this subset of plants we see similar trends through time: host identity effects diminish (Harvest 1: Adj. R^2^ = 0.114, p < 0.001; Harvest 2: Adj. R^2^ = 0.050, p < 0.001; Harvest 3: Adj. R^2^ = 0.024, p = 0.007) and neighbor identity effects increase (Harvest 1: Adj. R^2^ = −0.003, p = 0.325; Harvest 2: Adj. R^2^ = 0.010, p = 0.031; Harvest 3: Adj. R^2^ = 0.018, p = 0.013). In this case a significant block effect was only observed at time point 3 (Adj. R^2^ = 0.032, p = 0.013), suggesting the block effect trend described above is influenced by the no neighbor controls.

We further asked whether a closer approximation of bacterial taxon absolute abundances might impact our conclusions. The relative abundance of each taxon in each sample was multiplied by the total 16S rRNA copies per 10 μl DNA (the same volume used for sequencing library preparation) to yield an estimate of each taxon’s absolute abundance (quasi-absolute abundance). In this new dataset, sample dissimilarities were modeled using the same PERMANOVA procedure as above. This generated the same qualitative findings as the relative abundance data, but with slightly stronger effect sizes for influence of neighborhood plant species (Supp. Table 1). One new result revealed by this approach, however, was a host by neighborhood interaction, which increased from harvest 2 (Adj. R^2^ = 0.008 p = 0.002) to harvest 3 (Adj. R^2^ = 0.024, p = 0.003, see Supp. Table 1 for pre-adjusted R^2^ values). In other words, the effect of neighborhood depended on the host’s species identity, and this effect became stronger over time.

To further assess whether the three host plants species differed in their susceptibility to neighborhood effects, we subset the data by plant species and assessed the strength of neighborhood effects separately over the 3 time points. Similar to the combined data, no plant species exhibited a detectable neighborhood effect of microbiome composition at harvest 1. The tomato focal plants only exhibited a detectable neighborhood effect at harvest 3 (R^2^ = 0.086, P = 0.009, Fig. 3B), and no block effects at any harvest. For the pepper focal plants, there was only a neighborhood effect at harvest 2 (R^2^_adj_ = 0.108, P = 0.013, Fig. 3B). Lastly for the bean focal plants, a neighborhood effect was only detected at harvest 3 (R^2^ = 0.069, P = 0.012, Fig. 3B). No significant block effects were observed for pepper or bean at any harvest.

#### Neighbor status and identity diminish effects of host species identity on phyllosphere bacterial communities

To test the hypothesis that focal plants with experimental neighbors would experience higher rates of inter-host dispersal than focal plants without nearby neighbors, we performed model selection on hosts that were distinguished by their neighbor status to identify the most explanatory host variables. We repeated this procedure for spatial variables generated by principle coordinates of neighborhood matrix analysis and then performed variation partitioning to partition out the unique contribution of host factors explaining diversity. We find supporting evidence for our hypothesis in harvests 2 and 3, but not harvest 1 (Fig. 4), indicating a dependency on the age structure of neighbors. For harvests 2 and 3, the unique contribution of host identity was stronger for the plants having no neighbors than the plants having either tomato, pepper, or bean as neighbors. In fact, at harvest 3 focal plants that were surrounded by bean or tomato neighbors had no detectable effect of host species identity after separating out the effect of space. Interestingly at harvest 1, focal plants with neighbors had higher effects of host species identity than the no neighbor controls, and this was especially the case for plants with tomato or bean neighbors. Interestingly in the focal plants surrounded by peppers, host species effects followed a hump-shaped relationship (increasing at first, then decreasing), rather than the monotonic decrease observed for tomato- or bean-surrounded plants. It thus appears that host-mediated selection of epiphytic bacterial communities becomes subservient to the effect of immigrant inoculum from neighboring plants as the biomass of neighboring plants increases.

**Fig. 4:**
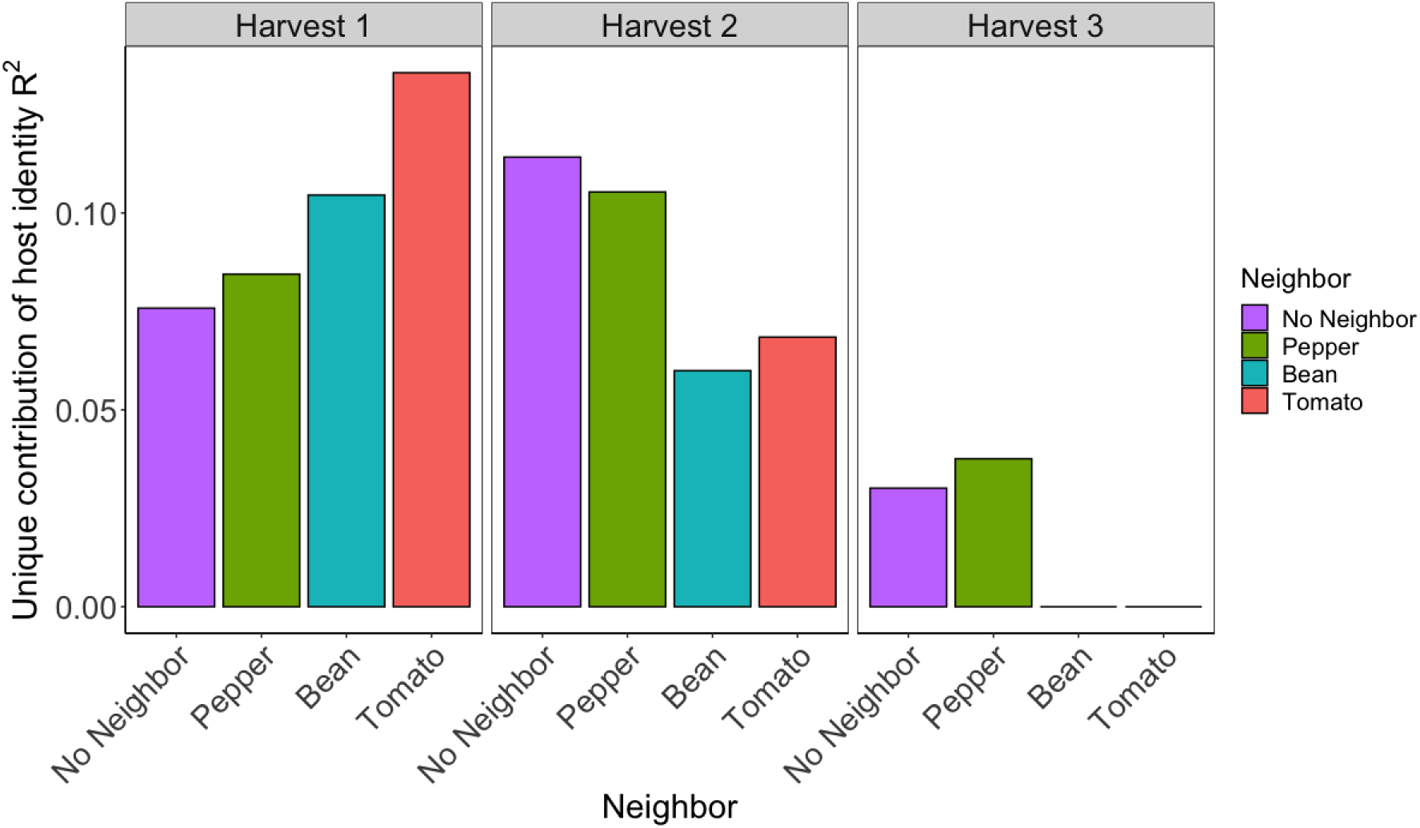
The unique contribution (adjusted R^2^) of host plant identity to phyllosphere bacterial structure decreases over time for plants with neighbors. Unique contribution was calculated by partitioning out spatial principle coordinates using RDA-based variation partitioning. The order of depiction of the neighbor plant species is by estimated neighborhood biomass from lowest to highest. Boxes 1, 2, and 3 represent different harvest time points. In cases where R_adj_ = 0, host species identity did not significantly explain variation in phyllosphere bacterial composition.

Additionally, in certain cases, host identity combined with host height, weight, or both height and weight in a way that increased explanatory power of host factors. While in several instances this boosted the explanatory power of host factors (e.g. at harvest 1), our qualitative conclusions remain the same (Supp. Fig. 1). In other words, the effects of host identity are weaker for all plants that have neighbors at both harvests 2 and 3.

#### Phylogenetic clustering and neutral model fit vary by host and through time

To better understand the predominance of deterministic processes in shaping phyllosphere community membership and determine whether the three plant species might be influenced by different assembly processes, we tested for patterns of phylogenetic clustering. Evidence of phylogenetic clustering within a host species would suggest that phylogenetically-conserved traits are being selected for in a host-specific way ^19,20^. We tested this idea using the standardized effect size (SES) of the mean pairwise distance (MPD) of bacterial ASVs in each sample. MPD SES was significantly influenced by host species identity (F_2,175_ = 219.86, p < 0.001), harvest time point (F_2,175_ = 12.33, p < 0.001), a host by harvest interaction (F_4,175_ = 43.84, p < 0.001), and neighborhood (F_3,175_ = 3.38, p < 0.05, Fig. 5A). Tomato- and pepper-associated communities were more phylogenetically clustered than would be expected by chance (as indicated by negative SES values), and this was the case for all three time points for both plants. In contrast, bean-associated communities showed evidence of phylogenetic overdispersion (as indicated by highly positive SES values). Bean SES values tended to decrease (i.e. tend towards clustering) over time, but were highly variable. One exception was at harvest 3, where beans with no neighbors had high variability in MPD SES, beans with conspecific neighbors were phylogenetically clustered, and beans with tomato neighbors were overdispersed (Tukey’s HSD conspecific neighbor vs tomato neighbor p = 0.02).

**Fig 5:**
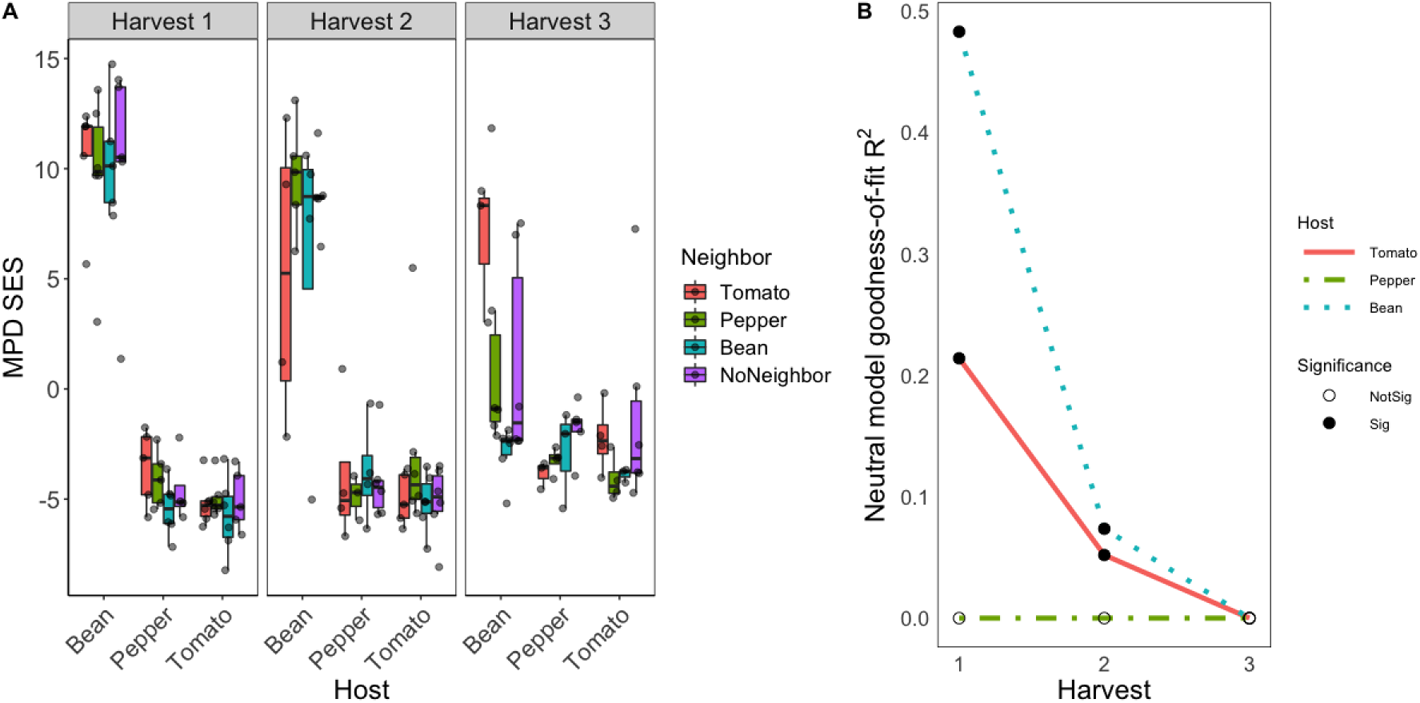
Bacterial leaf surface community assembly processes differ between plant hosts, suggesting differences in host filtering. A) Standardized effect size (SES) of mean pairwise distance (MPD) of phyllosphere communities split by host (x-axis), neighbor (box color), and harvest time point (panels 1, 2, or 3). SES = (MPD_obs_ –MPD_null_)/SD(MPD_null_), whereby values below 0 suggest phylogenetic clustering. B) The fit of a neutral model declines through time, but differs strongly by host identity. Neutral model goodness-of-fit values (R^2^, y-axis) at each harvest (x-axis) for tomato (solid red line), pepper (green dot-dashed line), and bean (blue dotted line). Filled circles indicate statistical significance, open circles indicate not significant (negative or 0 goodness-of-fit values).

We next asked how well the occupancy-abundance relationships within each host species could be fit by a neutral model, whereby passive dispersal and ecological drift are the primary drivers of establishment, and then asked whether the fit to that model changed over time. Of the three hosts, bean-associated communities had the highest goodness-of-fit values followed by tomato-associated communities, suggesting differences among hosts in the role of neutral processes in shaping community structure (Fig 5B). Both bean and tomato hosts showed a decline in the fit of a neutral model from harvest 1 to 2 (Bean: harvest 1 R^2^=0.483, harvest 2 R^2^=0.074; Tomato: harvest 1 R^2^=0.214, harvest 2 R^2^=0.052), and the neutral model failed to fit either set of plants for harvest 3 (as indicated by a negative goodness-of-fit). At all three time points, pepper hosts were never fit by a neutral model (indicated by negative goodness-of-fit values).

#### Experimental greenhouse transplantation of field study-derived inocula replicates host filtering and reveals effects of inoculum source

The subsequent greenhouse study allowed us to more closely examine the effects of inter-host dispersal of bacterial taxa, as phyllosphere bacterial communities that were recovered from the field were transplanted onto either the same plant species from which they were collected, or onto the plant species that had previously surrounded the source plant. From the 105 total such reciprocal inoculations, 103 samples yielded sufficient high quality sequences for analysis. The resultant dataset contained 2,640,588 observations of 1734 ASVs, 1379 of which had greater than 10 observations, and 621 of which had over 100 observations.

We observed a linear relationship between the ASV-level richness of the sample from which the inoculum was derived and the number of inoculum ASVs that were detectable in the greenhouse samples (R^2^ = 0.105, p = 0.004, Supp. Fig 3). The number of overlapping ASVs between the inoculum and the experimental plants was significantly related to the host species identity (F_2,88_ = 8.503, p < 0.001), the previous host species from which the inoculum was sampled (F_2,88_ =3.871, p =0.028), and interactions between the host and previous host (F_4,88_ = 2.598, p=0.049) as well as between the host and previous neighbor (F_4,88_ =2.968 p=0.03). Similar to the field study, phyllosphere communities were dominated by the phyla Proteobacteria and Firmicutes. The most abundant bacterial orders were the Bacillales, Burkholderiales, Enterobacterales, and Pseudomonadales (Supp. Fig. 2). The bacterial community structure on treated plants differed significantly from that of control plants to which only sterile buffer had been applied (PERMANOVA R^2^ = 0.0156, p = 0.048).

We used the heat-killed inoculum as a control to gain insights into the level of host selection and subsequent inoculum establishment across our experimental plants. To do so, we asked how strongly phyllosphere microbiomes were differentiated by host species identity. Plants that received a heat-killed inoculum were less differentiable than plants that received a live inoculum (PERMANOVA Adj. R^2^ = 0.100, p = 0.04 versus Adj. R^2^ = 0.103, p < 0.001, respectively). The plants that received the sterile buffer as a control were even less differentiable by host species identity (Adj. R^2^ = 0.056, p = 0.07). Interestingly, if we subset samples based on whether experimental plants received an inoculum from heterospecific (different species) or conspecific (same species) hosts, we see that heterospecific transplants resulted in more differentiable hosts than conspecific transplants (Adj. R^2^ = 0.123, p < 0.001 vs Adj. R^2^ = 0.109, p < 0.001, respectively). It thus appears that the treatment plants receiving live cells more efficiently filtered communities, driving differentiation of host species, and that heterospecific inoculum sources further bolstered host differentiation.

Indicator taxon analysis allowed us to examine the taxa enriched on each of the three host species in the field trial (at harvest 2) and the greenhouse trial. Of the 34 taxa that distinguished pepper plants in the field, 6 were found in the greenhouse dataset (Supp. Table 2). The collective relative abundance of these taxa was significantly higher on pepper plants in the greenhouse than on tomatoes (Tukey’s HSD p < 0.001) or beans (Tukey’s HSD p < 0.001). Of the 65 taxa that distinguished tomato hosts in the field, 25 were detected in the greenhouse dataset (Supp. Table 2). The collective relative abundance of these taxa was significantly higher on tomato and pepper plants relative to beans (Tukey’s HSD p < 0.05 and p < 0.01, respectively). Of the 300 taxa that distinguished bean, 27 were found in the greenhouse (Supp. Table 2), but their collective relative abundance was not significantly different among hosts (p > 0.05).

#### The effect of donor plant biomass on recipient plant phyllosphere richness in the greenhouse depends on the origin of field inoculum

Within-host ASV-level richness of treatment plants ranged from 24 to 231 ASVs and varied significantly by host species identity (F_2,74_ = 12.41, p < 0.001), an interaction between host and previous host identities (F_4,74_ = 3.50, p < 0.05), and experimental block (F_4,74_ = 2.43, p = 0.058). Of the three plant species, peppers harbored significantly higher richness than tomatoes or beans (p < 0.001), which were indistinguishable from each other (p > 0.05). When the inoculation was a conspecific transfer (i.e. moving between two plants of the same species), a negative but weak relationship was observed between donor plant biomass and recipient plant richness (Adj. R^2^ = 0.09, p < 0.05, Fig. 6A). However, when the inoculation was a heterospecific transfer (i.e. between two different plant species), a positive and stronger relationship was observed between donor plant biomass and recipient plant richness (Adj. R^2^ = 0.21, p < 0.01, Fig. 6B). No significant differences in richness were observed between conspecific transplants and heterospecific transplants (p > 0.05).

**Fig 6:**
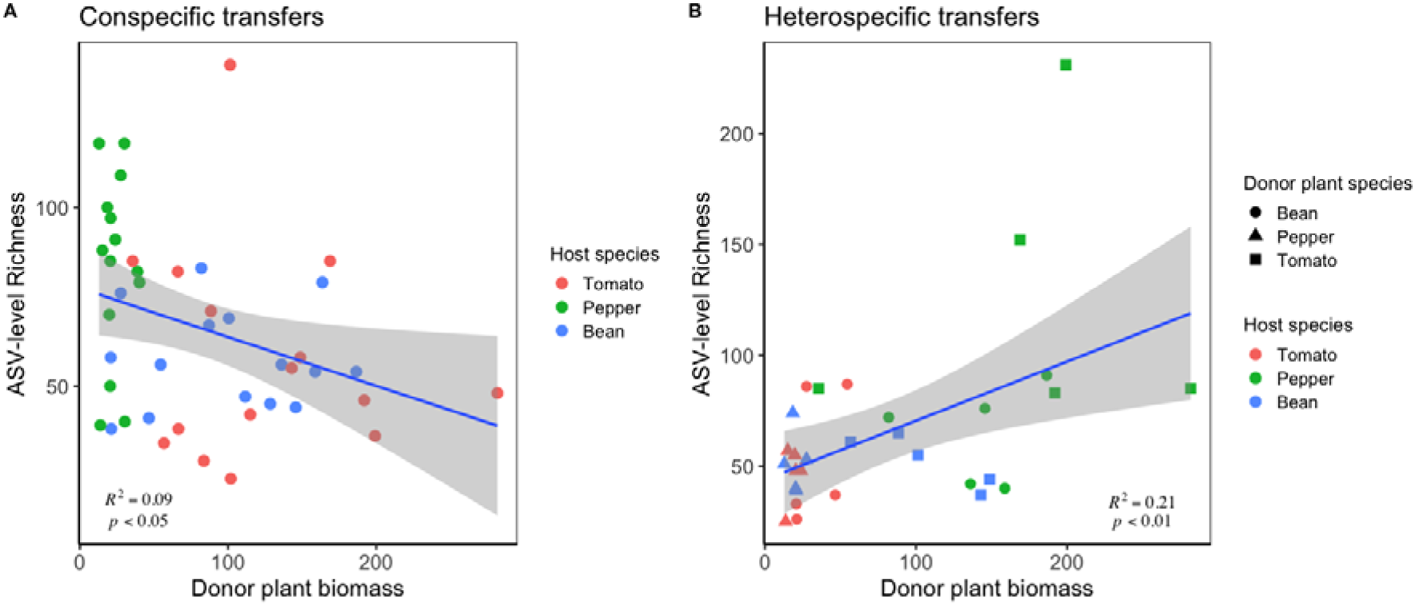
The effect of donor plant biomass (g) on phyllosphere community richness depends on the origin of inoculum. A) Conspecific (within the same species) transfers, where ASV-level richness (y-axis) is negatively, but weakly, correlated with the donor plant biomass (g, x-axis). Points are colored according to the recipient plant species. B) Heterospecific (across different species) transfers, where ASV-level richness (y-axis) is positively correlated with the donor plant biomass (g, x-axis). Point colors correspond to host species and point shapes correspond to the donor plant species. For both plots, R^2^ and p values are derived from linear models.

#### Experimental transplantation of phyllosphere communities reveals an influence of current and previous host identity, as well as donor plant neighbor

PERMANOVA on Bray Curtis dissimilarities revealed that host species identity, previous host identity (i.e. the species from which inoculum was derived), and experimental block all significantly contributed to phyllosphere community structure differences among experimental greenhouse plants (Host species R^2^ = 0.127, p <0.001, Previous host species R^2^ = 0.0468, p <0.001, Block R^2^ = 0.156, p <0.001, Table 2). We also observed an effect of donor plant biomass on recipient plant phyllosphere community structure (R^2^ = 0.018, p = 0.05, Table 2). Moreover, if we interrogate the dataset by plant species, we see a “grandparent effect” in the phyllosphere community structure of greenhouse-grown pepper plants. That is, pepper phyllosphere communities were most strongly shaped by the previous host’s neighbor (R^2^ = 0.194, p < 0.001), i.e. the plant species that previously surrounded the donor plant. The previous host’s species identity was also significantly associated with pepper plant phyllosphere community structure, albeit to a lesser extent (R^2^ = 0.107, p = 0.061). In contrast to what was observed on pepper, neither previous neighbor effects nor previous host effects were observed for tomato or bean subsets.

**Table 2:** Results of a PERMANOVA on phyllosphere bacterial community Bray Curtis dissimilarities for the greenhouse experiment. Variables tested include: host species identity (tomato, pepper, or bean), previous host species identity (tomato, pepper, bean), previous neighbor (tomato, pepper, bean), previous host biomass (g), and experimental block (1 through 5). R^2^ values represent the fit of the model.

| Greenhouse Trial |  |  |  |  |  |
| --- | --- | --- | --- | --- | --- |
|  | *df* | *Pseudo-f* | *R*<sup>2</sup> | *P* | *Significance* |
| Host Species | 2 | 6.21 | **0.127** | **0.001** | *** |
| Previous Host Species | 2 | 2.28 | **0.046** | **0.001** | *** |
| Previous Neighbor Species | 2 | 0.98 | 0.02 | 0.478 | NS |
| Previous Host Biomass | 1 | 1.72 | **0.018** | **0.05** | * |
| Block | 4 | 3.92 | **0.156** | **0.001** | *** |

## Discussion

Plant-microbe associations form in part through host filtering of microbiota that arrive via dispersal ^64^. Our study experimentally manipulated neighbor presence, identity, and age in order to understand how these factors influence host filtering of phyllosphere communities. Over the course of the experiment, we found that host species identity effects on focal plant microbiomes decreased, while the effects of neighborhood increased (Fig 3A). This finding builds on past studies showing that host species- or genotype-level differences in microbiota change over the growing season (e.g. Wagner et al. 2016; Chaparro, Badri, and Vivanco 2014; İnceoğlu et al. 2011). However, an important distinction is that we experimentally held host developmental stage constant throughout the experiment, thereby demonstrating that changes in host species identity effects over time are not simply due to host ontogeny, but hinge on characteristics of the neighboring plant community such as neighbor identity and biomass.

The increasing strength of neighborhood effects throughout the field experiment suggests that as neighboring plants grow and enrich for host species-specific microbes, they become larger sources of microbial propagules to their surrounding neighborhood, and thus alter the outcome of host filtering through compositional changes to the local species pool. Two other lines of evidence from our study further underscore the importance of neighbor identity for phyllosphere community assembly. First, when we directly control the directionality of dispersal in our greenhouse microbial transplant study, we see that the source of inoculum (i.e. the species identity of the donor plant) significantly contributed to the microbiome composition of recipient plants (Table 2). Second, the field experiment uncovered strong differences in host species identity effects depending on the identity of neighbors (Fig. 4). For instance at harvests 2 and 3, hosts that were surrounded by tomato or bean neighbors were substantially less differentiable in their phyllosphere community structure than plants surrounded by pepper or that had no neighbors. This suggests firstly that having a neighbor impacts the differentiation of hosts, but crucially that the outcome of neighborhood effects depends on neighbor identity. Interestingly, it has also been reported that inter-host dispersal among zebrafish greatly diminished genotype-level microbiome differences ^42^. Our results not only reinforce this concept in plants, but suggest that this effect depends on neighbor identity.

One of the key differences among experimental neighborhoods was plant biomass, which varied by species (Fig. 1C). This is likely an important driver of the neighborhood effects observed in this study. Larger biomass plants could harbor higher abundances of microorganisms by virtue of having more microbial habitat, thereby increasing the load of propagules dispersing onto focal plants. While we chose not to sample the epiphytic communities of the neighborhood plants in order to leave them undisturbed for the duration of the experiment, we observed a positive correlation between focal host biomass and epiphytic bacterial abundance. Thus, larger biomass neighborhood could have diminished the strength of host filtering through mass effects, i.e. rescuing via dispersal the taxa that went locally extinct due to host selection. We see evidence for the importance of neighbor biomass in several results. First, as the higher-biomass tomato and bean neighborhoods grew, we see that focal plant species identity effects became weaker, so much so that by harvest 3 hosts were indistinguishable by their species identity if they were surrounded by tomato or bean (Fig. 4). Interestingly, at harvest 1, relative to no neighbor controls, focal hosts surrounded by tomatoes or beans exhibited higher species identity effects, suggesting that at an early stage, neighbor plants may bolster host filtering by providing higher abundance and/or diversity of propagules. Moreover, the effects of the smaller pepper neighborhoods followed a similar but lagged trend, whereby host differentiation was highest at harvest time point 2, followed by more diminished host species effects at harvest 3. This could be driven by the observed slower growth of peppers relative to beans or tomatoes. In the greenhouse study, we see that the biomass of the donor plants significantly contributed to variation in both community composition (Table 2) and diversity (Fig. 6) of recipient plants. Here, the relationship between recipient plant phyllosphere richness and donor plant biomass was negative and weak if the transplant was conspecific (same species) but stronger and positive if it was heterospecific (between different species). Together these results underscore that biomass is not only an important component of neighborhood effects, but that such influences of biomass may depend at least partly on the identity of the neighbors.

Our study also makes clear that differences in the strength of host filtering across species may impact susceptibility to neighborhood effects. We found that plant species exhibited neighborhood effects differently through time (Fig. 3B). Together with the observation of a host-by-neighborhood interaction effect, these results suggest that the relative impact of local neighborhood differs among focal host species, perhaps due to differences in the degree to which dispersal versus host filtering influence phyllosphere assembly. For instance, host-specific carrying capacities could be driving the observed differences in bacterial abundances across species (Fig. 2A). This could mean that species with lower abundances of bacteria (e.g. peppers) are more invasible, and hence taxa that are selected for may more quickly become outnumbered by immigrating taxa. This may explain why pepper plants exhibited neighborhood effects at harvest 2, while tomato and beans did not do so until harvest 3. Several lines of evidence also suggest that the plants may differ in their selective abilities. For instance, tomato- and pepper-associated communities consistently exhibited phylogenetic clustering (Fig. 5A), indicating that closely related taxa were often observed to co-occur on a single plant, perhaps due to host selection of shared traits. In contrast, bean microbiomes tended to be phylogenetically overdispersed, and hence exhibited much less clustering than one would expect by chance. Overdispersion could indicate that traits under selection are not phylogenetically conserved, that competition is strong (e.g. between ecologically similar taxa), or that selection is relatively weak ^19^. That we also see a strong fit of bean microbiomes to neutral models at harvests 1 and 2 (Fig. 5B) suggests that the bean phyllosphere may be less selective relative to the tomato and pepper phyllosphere, and therefore more susceptible to dispersal effects.

Interestingly in the greenhouse experiment, only pepper plant microbiomes exhibited a “grandparent effect” of inoculum, i.e. an effect of the donor plants’ previous neighbor. This result demonstrates that for certain species, microbiome composition not only reflects its contemporary host and its source history, but also its previous dispersal history. This is analogous to a child inheriting a parent’s microbiome that carries with it traces of the parent’s former house, pet, or domestic partner. Results from the field trial may help shed light on why only pepper microbiomes contained detectable traces of dispersal history. The aforementioned patterns of phylogenetic clustering in pepper plant communities and the observation that neutral models failed to fit pepper plants (Fig. 5A,B) together suggest that the pepper phyllosphere may impose particularly strong selection on microbial communities. This strong filtering ability may not only select against taxa, but it could act to amplify taxa that previously dispersed from pepper neighbors, thus giving rise to effects of previous neighbor. In other words, selective plants such as pepper may bolster pepper-associated microorganisms upon arrival, even if they have become rare through multiple dispersal events.

One major implication of our work is that it highlights the potential importance of local host frequencies in recruiting microbiota. At harvest 3, bean phyllosphere communities exhibited phylogenetic clustering only when the plants were surrounded by conspecific (same species) neighbors. As phylogenetic clustering could be interpreted as host filtering for phylogenetically-conserved traits, this result suggests that for beans in particular, the frequency of conspecifics in the plant metacommunity may be an important determinant of host filtering efficacy. Interestingly, for pepper- or tomato-associated communities, the con- or hetero-specific status of neighborhoods had little influence over phylogenetic clustering, suggesting that this effect may depend on the strength of host filters. Similar findings have recently been reported for Acer saccharum trees ^38^, where the abundance of conspecific trees in the local metacommunity was positively correlated with the degree of host specialization in the phyllosphere, and importantly, this was not the case for all tree species surveyed. Moreover in cacao trees, leaf litter of healthy conspecific hosts was shown to protect against pathogen damage ^67^. While we did not test the fitness effects of host filtering for specialized microbial taxa, our results alongside several others may challenge the conspecific negative density-dependence (i.e. Janzen-Connell) hypothesis that posits that higher local densities of conspecifics may be disadvantageous due to the possibility of shared pests or pathogens ^29,30^. While there remain many examples of conspecific negative density-dependence ^68^, particularly in the tropics, meta-analyses seeking general trends have generated mixed results ^69,70^. Our work emphasizes that recruitment of specialized taxa from nearby conspecific hosts could outweigh the negative effects of pest pressure in certain contexts, or for certain host species.

Overall, our work makes clear that local neighborhood identity and biomass are key components that shape assembly of the phyllosphere microbiome. In both the field trial and greenhouse experiment, we find that although plants are able to select upon their microbial communities, the outcome of this selection is shaped by both neighbor identity and local biomass. Moving forward, this work has opened a number of critical questions regarding how neighborhood effects on the plant microbiome might shape plant health, fitness, and – in agricultural settings especially – yield. The work also raises questions about how invasive plant species might alter microbial dispersal within their communities, and potentially negatively feedback on native plant species’ fitness by reducing their ability to filter the optimal microbiome. In sum, our work demonstrates that host filtering and local dispersal are intimately intertwined and represent crucial considerations for the study of host-microbe associations.

## Acknowledgements

We acknowledge that this work was conducted on the territory of xučyun (Huichin), the ancestral and unceded land of the Chochenyo speaking Ohlone people; therefore this work would not have been possible were it not for their past land stewardship. We thank the staff of UC-Berkeley’s Oxford Tract Greenhouse for their assistance in maintaining the experiment. Thank you to R. Koutsoukis, T. Caro, X. Zhang, R. Debray, C. Hernandez, and E. Mehlferber for assistance with initial planting, and to the Koskella lab for invaluable feedback throughout.

