## Supplementary figures and images for "Plant neighborhood shapes diversity and reduces interspecific variation of the phyllosphere microbiome"

### Supplementary Figure 1

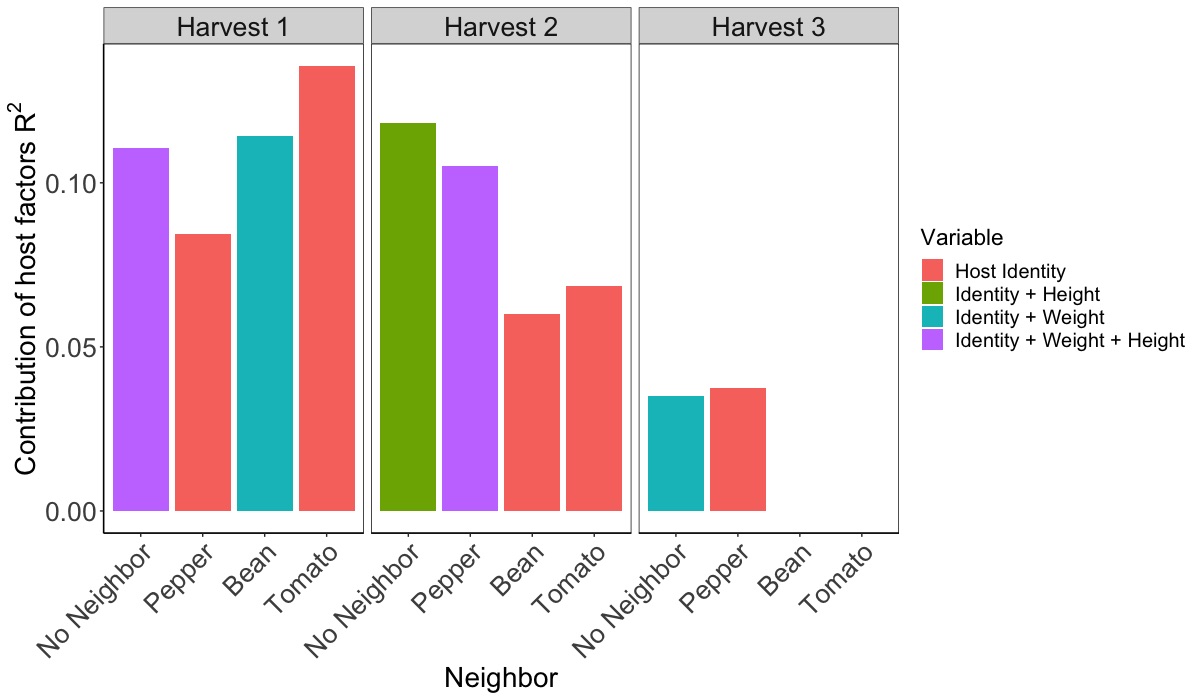

### Supplementary Figure 2

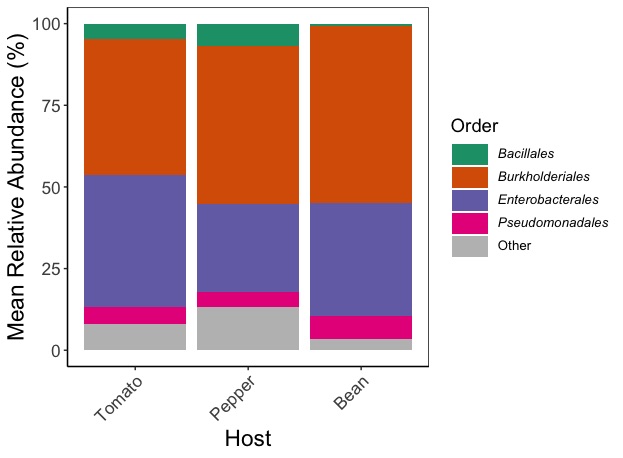

### Supplementary Figure 3

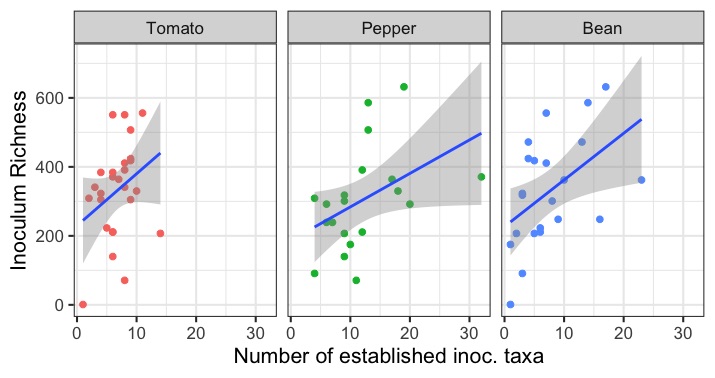
